# Paleozoic Protein Fossils Illuminate the Evolution of Vertebrate Genomes and Transposable Elements

**DOI:** 10.1101/2021.11.26.470093

**Authors:** Martin C. Frith

## Abstract

Genomes hold a treasure trove of protein fossils: fragments of formerly protein-coding DNA, which mainly come from transposable elements (TEs) or host genes. These fossils reveal ancient evolution of TEs and genomes, and many fossils have been exapted to perform diverse functions important for the host’s fitness. However, old and highly-degraded fossils are hard to identify, standard methods (e.g. BLAST) are not optimized for this task, and few Paleozoic protein fossils have been found.

Here, a recently optimized method is used to find protein fossils in vertebrate genomes. It finds Paleozoic fossils predating the amphibian/amniote divergence from most major TE categories, including virus-related Polinton and Gypsy elements. It finds 10 fossils in the human genome (8 from TEs and 2 from host genes) that predate the last common ancestor of all jawed vertebrates, probably from the Ordovician period. It also finds types of transposon and retrotransposon not found in human before. These fossils have extreme sequence conservation, indicating exaptation: some have evidence of gene-regulatory function, and they tend to lie nearest to developmental genes. Some ancient fossils suggest “genome tectonics”, where two fragments of one TE have drifted apart by up to megabases, possibly explaining gene deserts and large introns.

This paints a picture of great TE diversity in our aquatic ancestors, with patchy TE inheritance by later vertebrates, producing new genes and regulatory elements on the way. Host-gene fossils too have contributed anciently-conserved DNA segments. This paves the way to further studies of ancient protein fossils.

## Introduction

Genomes contain relics of formerly protein-coding DNA, which may be functionless and neutrally evolving, or in some cases have gained new, non-protein-coding functions. Most of them are derived from either transposable elements or host genes.

Transposable elements (TEs) are parasitic, or perhaps symbiotic, DNA elements that get copied or moved from one genome location to another. They have often proliferated greatly, so that e.g. the human genome has millions of TE-derived segments comprising at least ∼50% of the genome. Most of these segments are highly-mutated fragments, no longer active TEs.

TEs have had a massive impact on the evolution of their hosts (Warren et al. 2015; Etchegaray et al. 2021). They cause mutations by their proliferation, and also by ectopic recombination between TE copies, causing deletions, inversions, and duplications. This can duplicate or inactivate genes (Barsh et al. 1983; Hayakawa et al. 2001), or change their tissue-specific expression (Ting et al. 1992). Some host genes have evolved from TEs, such as the vertebrate RAG genes that generate the diverse antibodies and T cell receptors of the immune system (Kapitonov and Koonin 2015), and syncytin genes that seem to enable cell fusion in placental development (Dupressoir et al. 2005). Some DNA elements that regulate gene expression have also evolved from TEs (Ting et al. 1992; Jordan et al. 2003).

A series of studies in 2006–2007 found thousands of TE-derived non-protein-coding elements with strong evolutionary conservation in mammals (Bejerano et al. 2006; Nishihara et al. 2006; Xie et al. 2006; Kamal et al. 2006; Lowe et al. 2007; Gentles et al. 2007). They often occur in gene deserts, and nearest to developmental genes (Lowe et al. 2007). These TE insertions often predate the placental/marsupial divergence (Mesozoic), but few clearly predate the mammal/bird divergence (Paleozoic), and an exceptional handful (“at least several”) were shown to predate the amniote/amphibian divergence (Bejerano et al. 2006). It is thus remarkable that a later study claimed to find 133 TE insertions predating the divergence of humans and ray-finned fish, by comparing human TE fragments found by RepeatMasker to vertebrate genome alignments (Lowe and Haussler 2012).

The boundary between TEs and viruses is blurry, and an entire field, paleovirology, is mainly based on viral insertion fossils in eukaryote genomes (Barreat and Katzourakis 2021a). The oldest viral fossils found so far seem to be Mesozoic (Suh et al. 2014; Barreat and Katzourakis 2021a).

TEs are diverse and their classification is partly arbitrary (Storer et al. 2021; Kojima 2019), but eukaryotic TEs are conventionally split into *retrotransposons* which duplicate by reverse transcription of their RNA into DNA, and *DNA transposons* which do not. Major types of retrotransposon are: LINEs (“long interspersed nuclear elements”), LTR retrotransposons (which bear long terminal repeats), YR (tyrosine recombinase) retrotransposons, and Penelope-like elements. These are further subclassified, e.g. LINEs have clades and sub-clades such as Hero, Nimb, L1, I, and CR1. Major types of DNA transposon are: DDE transposons (named after 3 key amino acids in the transposase), Cryptons (YR transposons), Helitrons, and Polintons (also called Mavericks). These are also subdivided, e.g. DDE transposons have “superfamilies” such as Academ, hAT, Kolobok, and piggyBac. Finally, nonautonomous TEs such as SINEs (“short interspersed nuclear elements”) typically encode no proteins, and propagate by hijacking enzymes from autonomous TEs.

Many types of TE are patchily present and absent in closely- and distantly-related eukaryotes (Yuan and Wessler 2011; Chalopin et al. 2015). This can sometimes be explained by ordinary vertical inheritance, with multiple losses of the TE family (Fawcett and Innan 2016). Contrarily, it has been suggested that long-term vertical persistence of TEs may be rare, so their long-term persistence depends on horizontal transfer (Gilbert and Feschotte 2018). Thus, in order to understand the evolution of TE families in eukaryotes, it is valuable to know what TE types were present in ancestral eukaryotes (Fawcett and Innan 2016).

Host-gene-derived protein fossils are often called “pseudogenes”. They usually arise from duplication of (part of) a gene, such that one of the two copies is either not expressed or dispensable so evolves away from its protein-coding ancestry. Many such duplications are created by reversetranscription of mRNA to DNA (e.g. by retro-transposon enzymes), producing intron-depleted fossils termed “processed pseudogenes”. There are also non-duplicated “unitary pseudogenes”, e.g. the *GULO* /*GULOP* gene/pseudogene for making vitamin C, which is non-functional in primates and guinea pigs (Nishikimi et al. 1994).

Some pseudogenes seem to have significant functions, e.g. by being transcribed into an antisense RNA regulator of its cognate gene (Korneev et al. 1999), or regulating transcription (Huang et al. 2017), or generating small interfering RNAs (Tam et al. 2008). The Xist RNA involved in X chromosome inactivation has evolved partly from a formerly protein-coding gene, and partly from TEs (Elisaphenko et al. 2008). The boundary between protein fossils and functional protein-coding genes is fuzzy: they may produce peptides with marginal contribution to the organism’s fitness, in the process of gene death or resurrection (Brosius and Gould 1992; Cheetham et al. 2020).

Genetic fossils are often found by comparing a genome to a database of TE or gene sequences (Storer et al. 2021; Harrison 2021). This can be done by either DNA-to-DNA or DNA-to-protein comparison. Protein-coding DNA tends to evolve by changes that preserve the encoded amino acids or replace them with similar ones: thus highly-diverged sequences can be detected more effectively at the protein level (States et al. 1991). On the other hand, protein fossils evolve without amino-acid conservation. Thus, new TE families are often found by protein-level matches to distantly-related families, whereas relics of known TE families are best detected by DNA-level matches to a model approximating the family’s most-recent active ancestor. RepeatMasker files of such DNA-level matches are available for many genomes (Smit et al. 2015).

Protein-level matches have usually been sought with BLAST (Altschul et al. 1997), which is not optimized for fossils. Central to sequence matching methods are parameters defining the (dis)favorability of substitutions and gaps, which provide the definition of similarity. BLAST uses a 20×20 amino acid substitution matrix (BLOSUM or PAM), which is based on substitution rates in living proteins, so is likely suboptimal for fossils.

Therefore, we recently developed a new DNA-to-protein matching method (Yao and Frith 202x), implemented in LAST (https://gitlab.com/mcfrith/last). Its main advantage is that it sets the substitution, gap, and frameshift parameters by maximum-likelihood fit to given sequence data. It uses a richer 64×21 substitution matrix, allowing e.g. preferred matching of asparagine (encoded by `aac` or `aat`) to `agc` than to `tca`, which both encode serine. It judges homology based on not just one alignment, but on many alternative ways of aligning the putative homologs. This proved more sensitive than BLAST for finding human TE protein fossils, and for the first time it found YR retrotransposon fossils in the human genome (Yao and Frith 202x).

Here, this method is used to find new protein fossils in human and slowly-evolving Lagerstätte genomes: alligator, turtle, coelacanth (a lobe-finned fish closely related to land vertebrates), and chimaera (a non-bony cartilaginous fish); and also frog due to its intermediate phylogenetic position (table 1). The number of new fossils is relatively small, but they are especially ancient and include types of TE not found in human before. They thus illuminate the evolutionary history of TE content, and reveal strongly conserved ancient exaptations, including of host-gene fossils.

**Table 1.** Genome versions and TE protein fossils

| organism |  | genome assembly<br>(from NCBI or UCSC) | RepeatMasker<br>version (source) | TE<br>fossils | of which<br>novel* (%) |
| --- | --- | --- | --- | --- | --- |
| human | *Homo sapiens* | UCSC hg38.analysisSet | 4.0.7 (UCSC) | 546821 | 1641 (0.3) |
| alligator | *Alligator mississippiensis* | ASM28112v4 | 4.0.6 (NCBI) | 410092 | 46065 (11) |
| turtle | *Chrysemys picta bellii* | Chrysemys_picta_bellii-3.0.3 | 4.0.6 (NCBI) | 430459 | 63301 (15) |
| frog | *Xenopus tropicalis* | UCB_Xtro_10.0 | 4.0.8 (NCBI) | 135507 | 14837 (11) |
| coelacanth | *Latimeria chalumnae* | UCSC latCha1 | 4.0.5 (rmsk<sup>†</sup>) | 286944 | 279710 (97) |
| chimaera | *Callorhinchus milii* | UCSC calMil1 | 4.0.3 (UCSC) | 105995 | 31098 (29) |

\*Not found by this version of RepeatMasker
<sup>†</sup>repeatmasker.org

## Results and Discussion

### Protein-Fossil-Finding Pipeline

For each organism, homologous segments were found between the genome and a set of protein sequences comprising TE proteins from RepeatMasker plus proteins encoded by host genes of that organ-ism. When multiple homologies overlapped in the genome, only the strongest was kept, to avoid homologies between different types of TE or between TEs and host genes. Homologies overlapping annotated protein-coding segments of the genome were removed. Finally, host-gene homologies were discarded if they overlapped TEs annotated by RepeatMasker: this removes true-but-unwanted homologies due to host gene protein-coding segments that evolved from e.g. SINEs. The resulting fossils, including a genome browser hub, are available at https://github.com/mcfrith/protein-fossils.

The homology search used a significance threshold of one expected random match to the set of proteins per 10^9^ bp, so the human genome would have ∼3 matches if the sequences were perfectly random. The real false-positive rate will be higher, and was estimated by comparing the reversed (but not complemented) human genome to the proteins, producing 19 spurious matches (Yao and Frith 202x).

### New TE Fossils

The number of TE protein fossils found per genome ranges from ∼100 000 to ∼500 000, most of which correspond to known TE fragments in public RepeatMasker files (table 1). The human genome has especially few new TE fossils, indicating how thoroughly human TEs have been analyzed. The coelacanth fossils are almost all new relative to the RepeatMasker annotations, simply because those annotations have very few TE types, illustrating that TE analysis is lacking for some genomes at any snapshot in time (Sotero-Caio et al. 2017).

### Classifying Unknown Repeats

RepeatMasker genome annotations include repeats of unknown type, which might not be TEs. In alligator and turtle (but not the other genomes), some of these unknown repeats could be classified based on large and consistent overlaps with TE protein fossils (table 2). One of these repeats, UCON84, also occurs in the human genome: it is derived from a DDE transposon in the PIF/Harbinger superfamily (fig. 1). The UCON84 consensus sequence, obtained from Dfam (Storer et al. 2021), has shorter and weaker (but significant) homology to PIF/Harbinger proteins (not shown). The consensus is expected to approximate an ancestral sequence and thus have clearer homology, but it is hard to make an accurate consensus of ancient fragments.

**Table 2.** Classifying unknown repeats

| unknown repeat | TE type |
| --- | --- |
| REP-2_CPB | CR1 (LINE) |
| REP-3_CPB | L2 (LINE) |
| REP-6_CPB | CR1 (LINE) |
| REP-21_CPB | CR1 in PIF/Harbinger |
| REP-22_CPB | hAT-Tag1 |
| REP-28_CPB | CR1 (LINE) |
| REP-31_CPB | Gypsy (LTR) |
| SAT-928_Crp | Penelope |
| UCON84 | PIF/Harbinger |

**Fig. 1.**
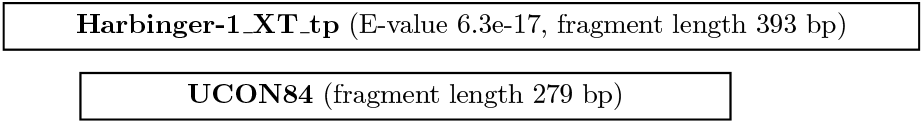
Overlap between a TE protein fossil (upper box) and a repeat of unknown type (lower box) in the alligator genome (at coordinate 15 466 729 in `NW_017707593.1`).

An interesting case is REP-21 CPB in turtle: its central region consistently overlaps CR1 LINE protein matches, while the left and right flanks consistently overlap PIF/Harbinger matches. Thus, it seems that a CR1 fragment was inserted in a PIF/Harbinger transposon, then this hybrid element proliferated, producing thousands of copies. We may speculate that it proliferated as a typical non-autonomous DNA transposon, with damaged protein-coding sequence but intact termini recognized by the transposase.

### Inter-Genome Homology

The age of genetic fossils can be inferred by comparing different genomes. For example, fig. 2 shows a human TE fossil aligned to an L1 LINE protein, alongside mammal genome alignments from the UCSC genome browser (Kent et al. 2002; Harris 2007). This L1 insertion is present in various monkey genomes but absent from bushbaby and other placental mammals, showing that the insertion occurred in a common ancestor of monkeys. It is thus curious that the L1 insert is aligned to two marsupial genomes, opossum and tasmanian devil. Marsupials also have L1s, and these marsupial regions are indeed annotated as L1s by RepeatMasker. Thus, these human and marsupial *inserts* are true homologs, because all L1s share common ancestry, but the *insertions* are not homologous: not descended from a common ancestral insertion. The inserts might even be orthologs, if their common ancestor is no older than the placental/marsupial divergence.

**Fig. 2.**
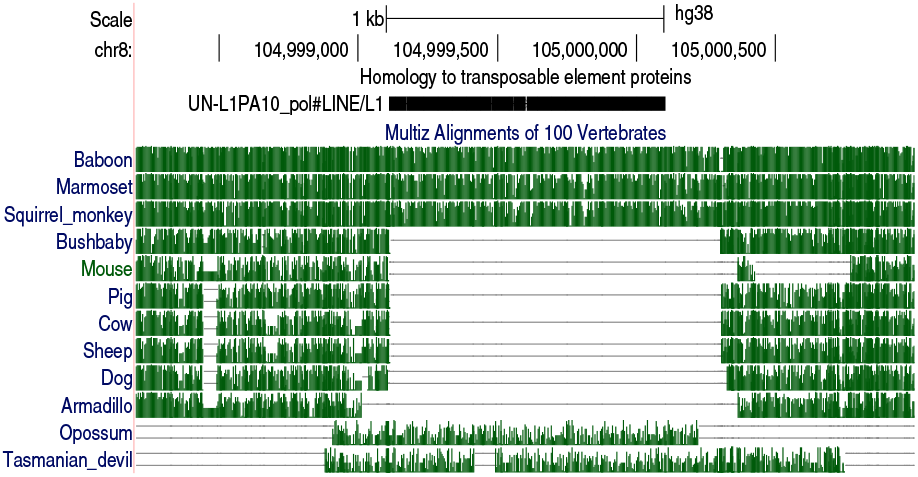
A TE protein fossil in human chromosome 8, with confusing inter-genome homology. Black bar near top: alignment of an L1 LINE protein. Green tracks: alignments between the human and other genomes. Screen shot from http://genome.ucsc.edu.

Why, then, do these marsupial alignments extend into flanking sequence beyond the insert? It is hard to determine the precise endpoint of homology between distantly-related sequences: alignments overshoot or undershoot. These human-marsupial alignments were made with the HoxD55 substitution matrix and gap parameters that are prone to large overshoots (Frith et al. 2008).

For this study, new pair-wise genome alignments were made, by finding homologous regions (Frith and Noée 2014) and cutting them down to most-similar one-to-one alignments (Frith and Kawaguchi 2015). This tends to find higher-similarity alignments than those from UCSC and elsewhere, indicating that a higher fraction of the alignments are orthologous (Frith and Kawaguchi 2015). This probably does not avoid non-homologous TE insertions, so a new step was added: isolated alignments were discarded, by only keeping groups of alignments that are nearby in both genomes. Some examples are in fig. 3: each panel shows one TE fossil in the human genome (central vertical stripe) that overlaps an inter-genome alignment (diagonal lines/dots). The alignments are not isolated: they are flanked by other alignments, indicating homology of not just the TE insert but also the flanking regions. Because these are distantly-related genomes, most of the DNA lacks similarity and is unaligned. The alignable fragments are probably conserved by natural selection.

**Fig. 3.**
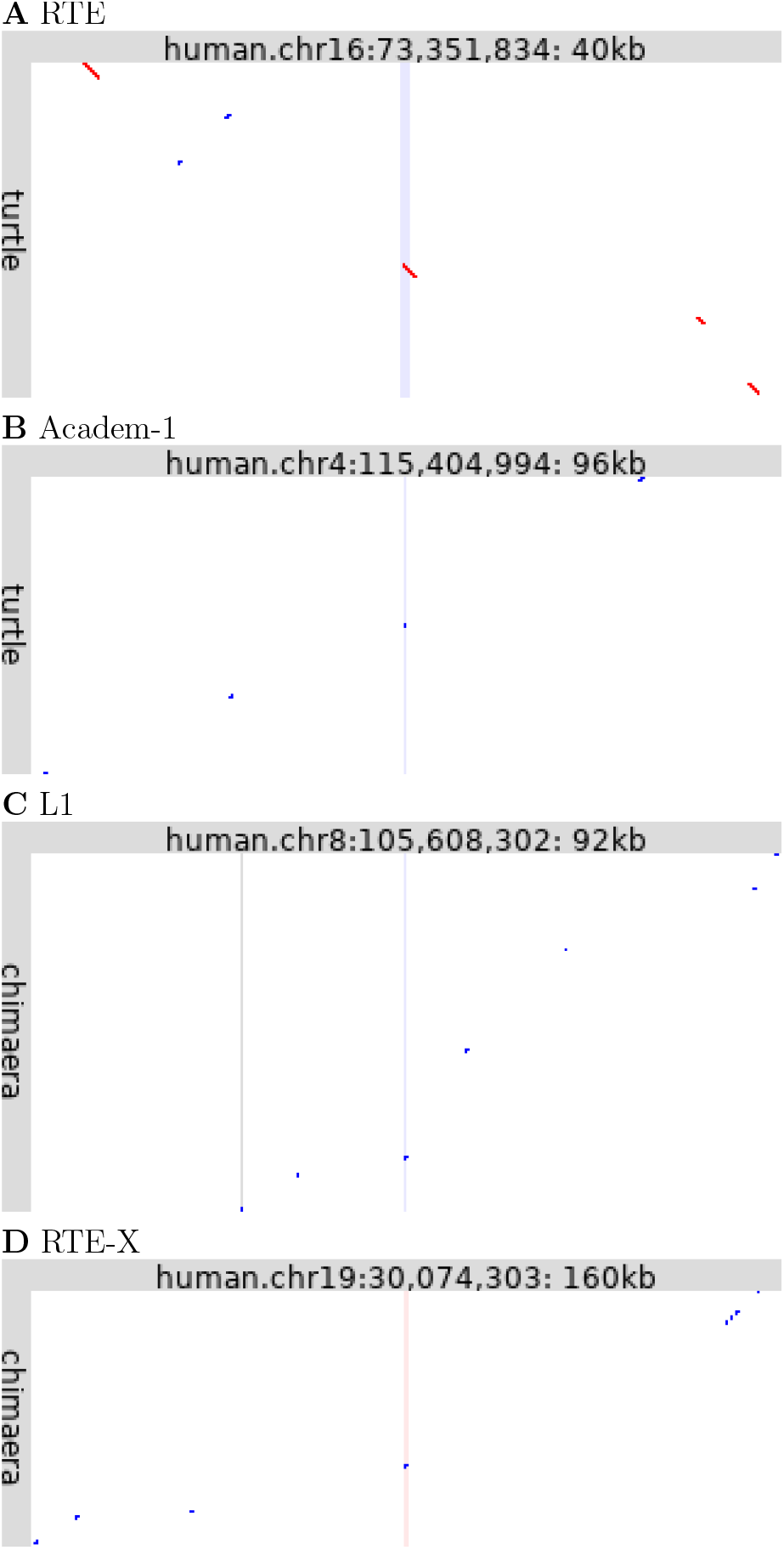
Ancient conserved TE insertions. Each panel shows alignments between part of the human genome (horizontal) and turtle (**A, B**) or chimaera (**C, D**). Red dots indicate same-strand alignments, blue dots opposite-strand alignments. The central vertical lines show the location in human of the TE fossil (pink: forward strand, blue: reverse strand). The vertical gray line in panel **C** shows a protein-coding exon of *ZFPM2*.

A possible objection is that these examples might be independent insertions of an abundant TE into homologous regions of two genomes. This cannot be ruled out, but the key point is that these alignments are not only homologies but most-similar one-to-one homologies: it would be a strong coincidence for these single-best matches to independently be in homologous regions.

### TE Types Newly Found in Human

The human TE protein fossils include several types of TE that have not been found in human before (table 3). These are all LINEs or DDE transposons, and are in addition to the first human YR retrotransposons (DIRS and Ngaro) and first-but-one Polintons we recently reported (Yao and Frith 202x). Some were found directly in human, others were found in another genome and mapped to human via the inter-genome alignments (“found in” column). The E-value indicates significance/confidence of the DNA-protein homology: it is the expected number of times to find such a similarity between the genome and the set of proteins, if they were random sequences. Some of the E-values are quite high, indicating lower confidence. On the other hand, most of these putative DNA-protein homologies overlap human/non-mammal genome alignments, which would be a strong coincidence if they were random similarities (fig. 3).

**Table 3.**
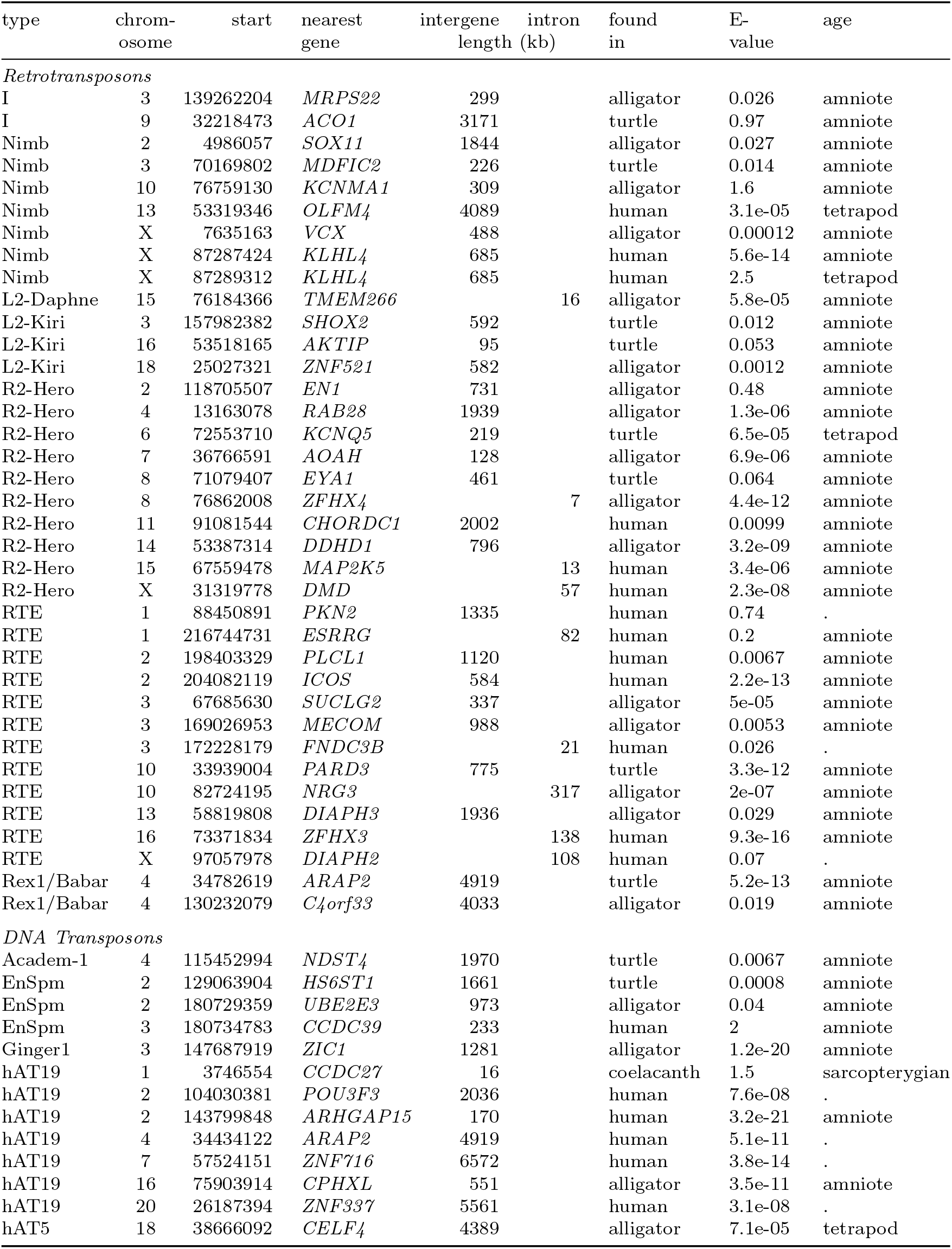
TE fossils of types newly found in human

Some of these TE fossils are shown in more detail in fig. 4. It can be seen that they lie in human genome regions conserved in non-mammals, and are not annotated by RepeatMasker. These regions have strong evolutionary conservation in mammals according to phastCons (Siepel et al. 2005), independent of their conservation in non-mammals. Some of these fossils overlap candidate regulatory elements or known transcription factor binding sites (Moore et al. 2020; Lesurf et al. 2016): the Hero fossil in fig. 4B overlaps a CEBPB binding site, and the RTE fossil in fig. 4C overlaps binding sites for GATA2, STAT1, JUND, FOS, and JUN. Fig. 4A shows two Nimb fragments that coincide with conserved DNA segments: presumably they come from one Nimb insertion, which predates the amniote/amphibian divergence. (Only one of these Nimb fragments is aligned to frog: the other may be deleted or not detected in frog.)

**Fig. 4.**
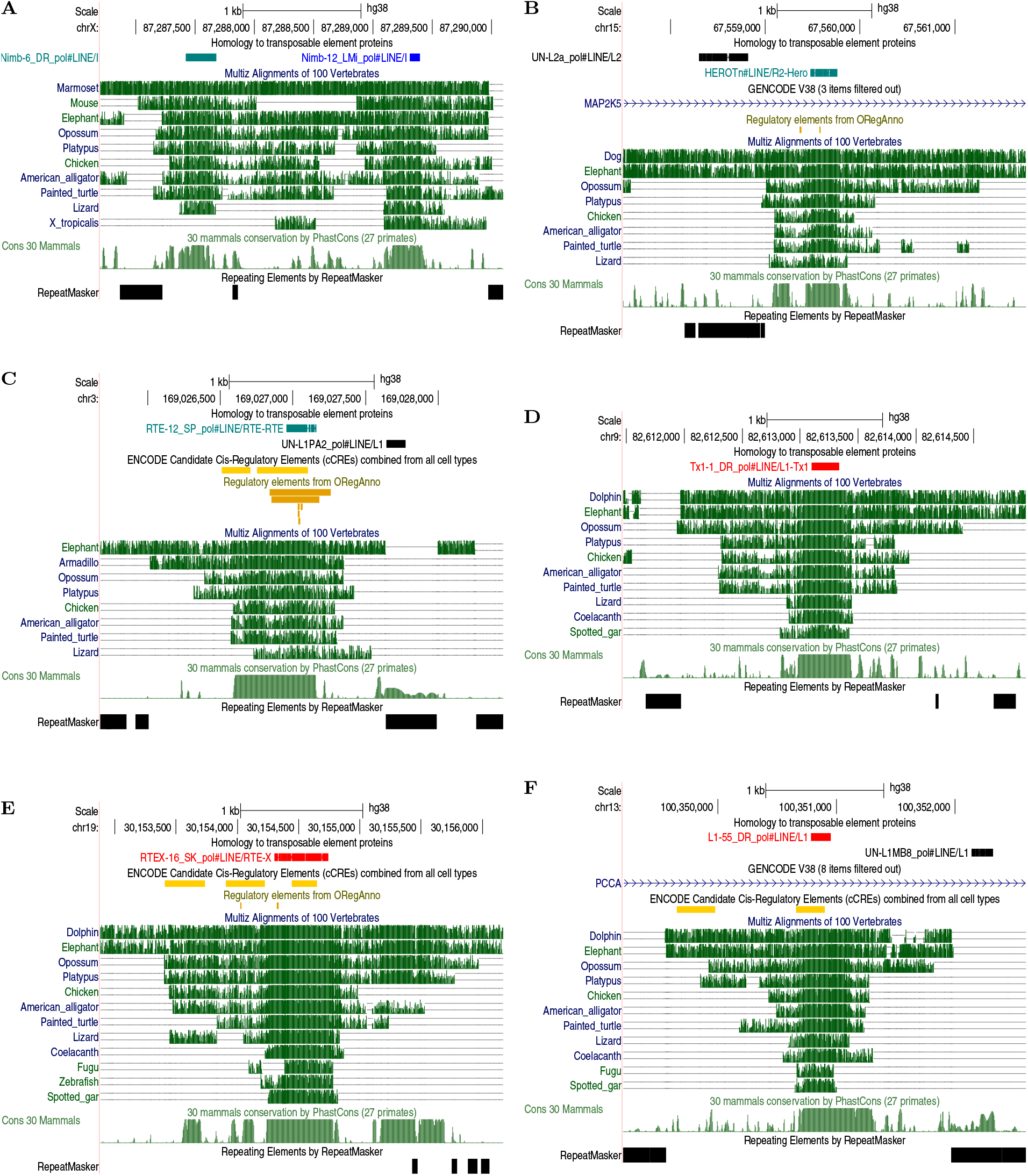
Ancient conserved TE insertions in the human genome. Each panel shows, from top to bottom: TE protein fossils, alignments of the human genome to other vertebrate genomes, evolutionary conservation in mammals (phastCons), and repeats found by RepeatMasker. Some panels also show annotations of regulatory elements and Gencode genes (introns). Screen shots from http://genome.ucsc.edu.

These fossils clarify the historical presence of TE types in vertebrates. They make presence of several TE types less patchy among vertebrates, thus explicable by vertical inheritance rather than horizontal transfer. Nimb-type LINEs have been found in insects, mollusks, teleost (bony) fish (Kapitonov et al. 2009; Chalopin et al. 2015), and turtle (Smit et al. 2015): here Nimb relics are found from ancient tetrapods, and also in coelacanth. This makes the presence of Nimb in vertebrates less patchy, and suggests vertical inheritance from the common ancestor of bony vertebrates. The Hero clade was found in sea urchin, lancelet, and fish (Kojima and Fujiwara 2004; Kapitonov et al. 2009): its presence in ancient tetrapods fits with vertical inheritance from deuterostome ancestors. Hero LINEs are unusual in having a restriction-like endonuclease (Kojima and Fujiwara 2004), unlike all other human LINE relics except Mam R4. The I clade was previously found in fish and some invertebrates (Kapitonov et al. 2009): here hundreds are found in turtle and a few in alligator. Daphne was previously found in sea urchin and arthropods (Schön and Arkhipova 2006), plus lancelet and zebrafish (Smit et al. 2015): here 67 fragments are found in coelacanth, 6 in chimaera, 6 in turtle, and 4 in alligator, rounding out its historical presence in vertebrates. The Rex1/Babar clade has been found patchily in non-sarcopterygian fish excluding chimaera, plus frog and lizard (Chalopin et al. 2015; Smit et al. 2015): here it is found in ancestral amniotes and also coelacanth and chimaera, rendering its distribution non-patchy.

RepeatMasker distinguishes two types of RTE-like LINE, BovB and RTE: it finds only BovB in human, and it finds RTE in turtle and zebrafish. Previous reports of RTE in human seem to be BovB elements that were not classified separately (Kojima 2018). This study finds RTEs from amniote ancestors, and thousands in coelacanth, again suggesting vertical inheritance from ancestors of bony vertebrates.

These fossils also provide support for TE origin of genes. Ginger1 transposons were previously found in some invertebrates including lancelet (Bao et al. 2010), but not in sarcopterygians (Yuan and Wessler 2011; Chalopin et al. 2015). Their relics are found here in alligator, turtle, coelacanth, and many in frog. This makes it more plausible that the human *GIN1* gene was indeed exapted from Ginger1 in ancestral amniotes (Bao et al. 2010). At least one pre-amniote Ginger1 relic was also exapted for non-protein-coding function (table 3). Similarly, hAT19 fossils from amniote ancestors support the hAT19 origin of the amniote-specific gene *CGGBP1* (Yellan et al. 2021), which binds CGG repeats and regulates gene expression (Singh and Westermark 2015). hAT19 fragments have been exapted for non-protein-coding functions too.

In the four tetrapod genomes just one hAT5 fragment is found, which is conserved in all of them: the single exapted relic of an ancient hAT5 infection. hAT5 was previously found in some invertebrates (Putnam et al. 2007) and fish (Smit et al. 2015), and is unusual in having 5 bp TSDs (target site duplications), whereas all previously-known hATs have 8 bp TSDs (Putnam et al. 2007).

### Anciently Conserved TE Fossils

The human genome contains diverse TE protein fossils that are older than the amniote/amphibian divergence (table 4). It is striking that they include nearly all major types of TE: LINEs, Penelope elements, LTR retrotransposons (Gypsy), YR retrotransposons (DIRS), DDE transposons, a Crypton, and Polintons. Eight of them (7 LINEs and a Crypton) are shared by human and chimaera, making them older than the last common ancestor of all jawed vertebrates. Three of these most ancient fossils are shown in fig. 4 D–F: their ancient exaptation is supported by their conserved presence in mammal, reptile, and bony-fish genomes, their strong conservation in mammals (phastCons), and sometimes by evidence of regulatory function.

**Table 4.** Pre-tetrapod TE fossils found in human

| type | chromosome | start | nearest gene | intergene length (kb) | intron | found in | E-value | age |
| --- | --- | --- | --- | --- | --- | --- | --- | --- |
| *Retrotransposons* |  |  |  |  |  |  |  |  |
| CR1 | 4 | 13161246 | *RAB28* | 1939 |  | alligator | 1.9e-06 | tetrapod |
| CR1 | 4 | 111235163 | *PITX2* | 1503 |  | chimaera | 0.0078 | gnathostome |
| CR1 | 5 | 94809895 | *MCTP1* |  | 69 | alligator | 0.0049 | tetrapod |
| CR1 | 6 | 98370405 | *POU3F2* | 1551 |  | turtle | 0.022 | tetrapod |
| CR1 | 16 | 78813330 | *WWOX* |  | 779 | turtle | 0.022 | tetrapod |
| CR1 | 18 | 72559688 | *CBLN2* | 198 |  | alligator | 9.9e-08 | tetrapod |
| Nimb | 13 | 53319346 | *OLFM4* | 4089 |  | human | 3.1e-05 | tetrapod |
| Nimb | X | 87289312 | *KLHL4* | 685 |  | human | 2.5 | tetrapod |
| L1 | 3 | 70416318 | *MDFIC2* | 642 |  | coelacanth | 0.61 | gnathostome |
| L1 | 8 | 105654302 | *ZFPM2* |  | 154 | human | 1 | gnathostome |
| L1 | 9 | 2050774 | *SMARCA2* |  | 7 | human | 0.093 | tetrapod |
| L1 | 13 | 100350784 | *PCCA* |  | 28 | human | 0.035 | gnathostome |
| L1 | 18 | 55784184 | *TCF4* | 961 |  | alligator | 6e-07 | tetrapod |
| L1-Tx1 | 9 | 82613098 | *RASEF* | 984 |  | human | 9.6e-05 | gnathostome |
| L1-Tx1 | 10 | 129519611 | *MGMT* |  | 69 | coelacanth | 3e-16 | gnathostome |
| L2 | 5 | 166711049 | *TENM2* | 3554 |  | turtle | 0.4 | tetrapod |
| L2 | 6 | 45881206 | *CLIC5* | 347 |  | alligator | 0.032 | tetrapod |
| L2* | 7 | 108869078 | *DNAJB9* | 2088 |  | human | 2.9e-25 | tetrapod |
| L2-Crack | 1 | 216137121 | *USH2A* |  | 77 | alligator | 3.5e-08 | tetrapod |
| L2-Crack | 5 | 109502097 | *PJA2* | 280 |  | human | 0.00013 | tetrapod |
| R2-Hero | 6 | 72553710 | *KCNQ5* | 219 |  | turtle | 6.5e-05 | tetrapod |
| RTE-X | 19 | 30154303 | *ZNF536* | 209 |  | human | 0.00078 | gnathostome |
| Penelope | 4 | 187183800 | *FAT1* | 1272 |  | alligator | 0.05 | tetrapod |
| Penelope | 13 | 106429488 | *EFNB2* | 999 |  | human | 0.00017 | tetrapod |
| Gypsy | 1 | 38669733 | *RRAGC* | 791 |  | human | 5.9e-11 | tetrapod |
| Gypsy | 2 | 57598618 | *VRK2* | 1521 |  | alligator | 8.3e-12 | tetrapod |
| Gypsy | 19 | 30111148 | *URI1* | 209 |  | human | 0.017 | tetrapod |
| Gypsy | X | 98972571 | *PCDH19* | 2687 |  | human | 4.6e-23 | tetrapod |
| DIRS | 3 | 55911374 | *ERC2* |  | 62 | alligator | 1.1e-08 | tetrapod |
| DIRS | 9 | 13728242 | *NFIB* | 802 |  | human | 0.0037 | tetrapod |
| DIRS | 9 | 20189423 | *MLLT3* | 553 |  | turtle | 0.86 | tetrapod |
| DIRS | 10 | 16734086 | *RSU1* |  | 57 | alligator | 0.84 | tetrapod |
| DIRS | 16 | 78152629 | *WWOX* |  | 49 | turtle | 3.2e-17 | tetrapod |
| *DNA Transposons* |  |  |  |  |  |  |  |  |
| PIF/Harbinger | 2 | 145279820 | *ZEB2* | 3324 |  | human | 4.4e-05 | tetrapod |
| PIF/Harbinger | 2 | 176676027 | *MTX2* | 875 |  | human | 0.0012 | tetrapod |
| hAT-Blackjack | 7 | 14272601 | *DGKB* |  | 160 | alligator | 6.5e-10 | tetrapod |
| hAT-Tip100 | 4 | 4696650 | *STX18* | 317 |  | alligator | 0.0021 | tetrapod |
| hAT-Tip100 | 4 | 129428560 | *C4orf33* | 4033 |  | human | 1.8e-11 | tetrapod |
| hAT19 | 1 | 3746554 | *CCDC27* | 16 |  | coelacanth | 1.5 | sarcopterygian |
| hAT5 | 18 | 38666092 | *CELF4* | 4389 |  | alligator | 7.1e-05 | tetrapod |
| Crypton-A | 12 | 14468609 | *ATF7IP* |  | 9 | alligator | 2.2e-70 | gnathostome |
| Polinton | 3 | 114555204 | *ZBTB20* |  | 83 | human | 2.8 | tetrapod |
| Polinton | 20 | 52577239 | *ZFP64* | 781 |  | turtle | 4.4e-15 | tetrapod |
| Polinton | 20 | 55516705 | *CBLN4* | 1346 |  | turtle | 7.4e-11 | tetrapod |

\*Previously found by RepeatMasker

A further 882 TE protein fossils that predate the mammal/reptile divergence were found in the human genome (table 5). Most of these (745, 84%) are novel (not annotated by RepeatMasker), as are all but one of the pre-tetrapod fossils (table 4). These ancient TE fossils are often in megabase-scale gene deserts or large (∼ 10^5^ bp) introns (table 3, 4). The nearest genes are significantly enriched in developmental functions such as nervous system development, cell morphogenesis, and axonogenesis (Mi et al. 2021).

**Table 5.** Other pre-amniote TE fossils in human

| type | number | of which known* |
| --- | --- | --- |
| *Retrotransposons* |  |  |
| CR1 | 204 | 92 |
| Dong-R4 | 1 | 0 |
| Vingi | 1 | 0 |
| L1 | 137 | 22 |
| L1-Tx1 | 12 | 0 |
| L2 | 167 | 14 |
| L2-Crack | 25 | 1 |
| BovB | 28 | 1 |
| RTE-X | 4 | 0 |
| Penelope | 54 | 1 |
| Gypsy | 83 | 3 |
| DIRS | 37 | 0 |
| Ngaro | 28 | 0 |
| *DNA Transposons* |  |  |
| Kolobok-T2 | 1 | 0 |
| PIF/Harbinger | 18 | 1 |
| PiggyBac | 7 | 0 |
| TcMar-Mariner | 1 | 0 |
| TcMar-Pogo | 2 | 0 |
| TcMar-Tc1 | 4 | 0 |
| TcMar-Tigger | 5 | 0 |
| hAT-Ac | 7 | 1 |
| hAT-Blackjack | 12 | 1 |
| hAT-Charlie | 6 | 0 |
| hAT-Tip100 | 21 | 0 |
| Crypton-A | 3 | 0 |
| Polinton | 14 | 0 |

\*Previously found by RepeatMasker

These TE relics from ancient vertebrates help us to understand the ancestral mobilome, which has been difficult, especially since TEs might have been horizontally transferred (Chalopin et al. 2015). For example, it has been suggested that mammal L1s were introduced by horizontal transfer into a common ancestor of therian (live-bearing) mammals (Ivancevic et al. 2018). We now have direct evidence that L1-like TEs were present in a common ancestor of jawed vertebrates, and hundreds of L1 fragments predate the amniote divergence (table 5). Actually, repeatmasker.org lists 353 L1 fragments in the platypus genome (`ornAna1`), so perhaps L1s were vertically inherited by mammals, but became inactive early in the monotreme lineage. There are also L1-Tx1 fossils from gnathostome ancestors (table 4): this supports the suggestion that L1 clades including Tx1 diverged in a common ancestor of mammals and fish (Ichiyanagi et al. 2007), which was not certain since Tx-like L1s are prone to horizontal transfer between marine hosts (Ivancevic et al. 2018).

For other TE types too – DIRS, Polinton, and PIF/Harbinger – their previously-noted patchiness among tetrapods (Chalopin et al. 2015) is explained by ancient loss of activity, since they were present in tetrapod ancestors. The emerging picture is that ancient vertebrates had many diverse types of TE, like present day teleost fish but unlike mammals or birds (Chalopin et al. 2015).

The pre-amniote BovB fossils (table 5) are particularly informative, because BovB has frequently been horizontally transferred (Ivancevic et al. 2018). Interestingly, the phylogeny of BovB elements differs greatly *but not entirely* from the phylogeny of their host organisms: amniote BovBs are all in a central branch of the tree and fish BovBs on outer branches (Ivancevic et al. 2018, fig. 2a). Knowing that BovBs were present in amniote ancestors, it seems likely that BovB initially entered amniotes by vertical inheritance, perhaps specifically into squamate reptiles, before being horizontally transferred among amniotes and arthropod vectors.

Regarding LTR retrotransposons, it is intriguing that ancient Gypsy-like fossils are found (table 4, 5), but ancient ERV (endogenous retrovirus) fossils are not. The origin of vertebrate retroviruses has been debated (Hayward 2017): ERVs may have evolved from Gypsy-like elements in a common ancestor of amniotes and amphibians (Hellsten et al. 2010).

The age of the oldest Polinton insertions is greatly increased from 95 million years (Barreat and Katzourakis 2021b) to ∼350 million years (the amniote/amphibian divergence). It is possible, however, that these protein fossils are much younger than their insertions, if the intact TE benefits host fitness so remains intact (i.e. protein coding) by natural selection of the host. Intact TEs are usually thought not to benefit host fitness, but intact Polintons might protect the host from viruses, in particular iridoviruses that infect cold-blooded vertebrates (Barreat and Katzourakis 2021b). Nevertheless, the human Polinton relics are no longer intact, yet some have strong phastCons conservation in mammals indicating exaptation.

### Conserved RepeatMasker Fossils

For sake of comparison, the age of previously known TE fossils (from RepeatMasker) was inferred in the same way. RepeatMasker includes many more TE fossils, especially non-protein-coding SINEs. It is tuned to have a false positive fraction of 0.2% (Hubley et al. 2016), which corresponds to ∼10^4^false hits in the human genome. There are 133 RepeatMasker hits in human that are conserved in frog, of which 84 (63%) are especially ancient types of repeat: UCON, Eulor, LFSINE, and AmnSINE1 (Gentles et al. 2007; Bejerano et al. 2006; Nishihara et al. 2006). Most of these are unknown types of repeat, and may not be TEs. In contrast, there are 73 RepeatMasker hits in human that are conserved in coelacanth, which are not obviously enriched in ancient repeat types. They include primate-specific L1P and SVA elements, which are surely false-positive RepeatMasker annotations. A few may be real, but it is hard to know which ones or have confidence in them. Unfortunately, RepeatMasker files do not state the significance (E-value) of each hit. In summary, the oldest confident minimum age for previously-known TE insertions (apart from TE-derived genes) is the amniote/amphibian divergence (Bejerano et al. 2006).

This casts doubt on the previously reported TE insertions predating the human/teleost divergence (Lowe and Haussler 2012). Aside from false Repeat-Masker hits, that study mentioned no countermeasures for non-homologous insertions (fig. 2).

The tetrapod TEs found here (table 4) are almost completely disjoint from previously known ones: the latter are mostly unknown repeat types or SINEs. The newly-found LINEs might be the autonomous counterparts of the ancient SINEs, in particular, AmnSINE1 was thought to be mobilized by an undiscovered L2-like LINE (Nishihara et al. 2006).

### Genome Tectonics

Sometimes, two TE fossils of the same type lie strikingly near each other in the human genome. An example is in fig. 5: two Polinton relics are separated by 44 kb, which is remarkably close considering there are only 40 Polinton fragments in the genome. They might come from two independent insertions into a Polinton hotspot, but a simpler explanation is that they come from one Polinton, and drifted apart due to younger TE insertions between them. It is well known that old TEs get fragmented by younger insertions, but it is interesting to consider how far apart they can drift. If there is a locally higher rate of insertion than deletion, this might over time produce large introns and gene deserts. Ancient fossils can be markers of such long term rifting. Among the pre-amniote TE fossils, there are a few hundred such pairs separated by 30–3000 kb.

**Fig. 5.**
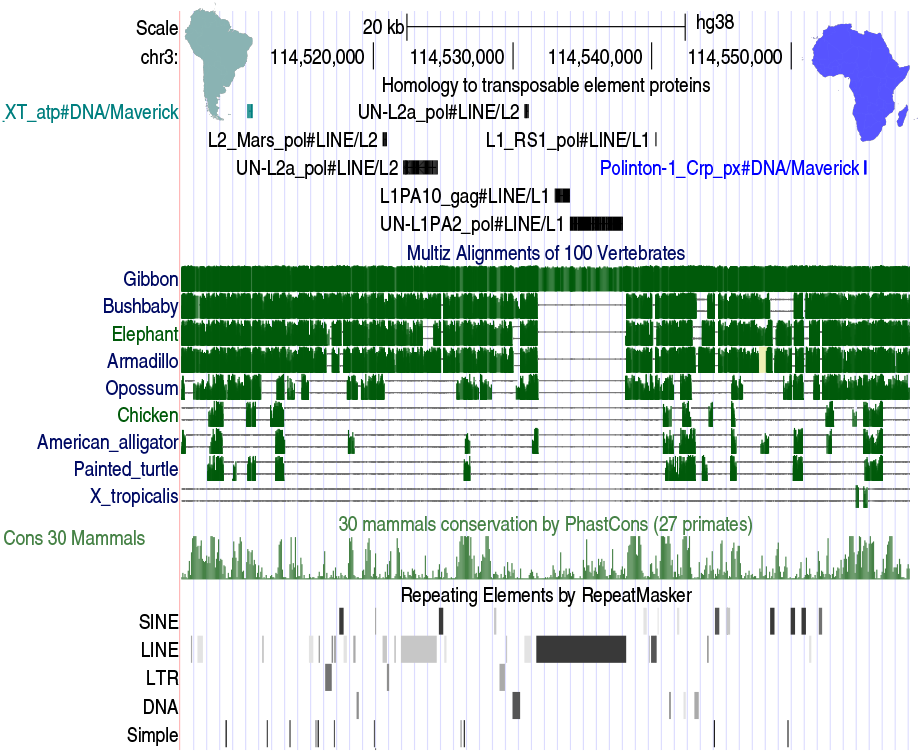
Ancient Polinton/Maverick fragments in an intron of *ZBTB20* on human chromosome 3. The two fragments are labeled by South America and Africa, with younger TE fossils in between.

### Host-Gene-Derived Protein Fossils

This study found 27 240 host-gene-derived protein fossils in the human genome, of which 4303 (16%) are new: not in Gencode V37 or RefSeq pseudogenes, or RetroGenes V9 (Pruitt et al. 2014; Harrow et al. 2012; Baertsch et al. 2008). They do not overlap known protein-coding regions, but some may be unknown protein-coding exons rather than fossils. Frameshifts or premature stop codons are present in 71.3% of the new segments and 72.4% of the non-new ones, suggesting a similar (presumably low) fraction of unknown coding exons.

Ancient fossils were sought in the same way as for TEs, but there is an extra difficulty. While we may find a fossil in the human genome that overlaps an alignment to (say) chimaera, it might have encoded a functional protein for most of this evolutionary history, becoming a fossil only recently in the human lineage (Sheetlin et al. 2014). The aligned region of chimaera was also required to be noncoding, but it may have independently become a fossil, or simply be an unannotated protein-coding exon. The non-human genomes presumably have less thorough gene annotation. This was assessed by manually examining UCSC phyloP graphs showing basewise evolutionary conservation in 100 vertebrates (Pollard et al. 2010). In some cases, there was a pattern of every 3rd base being less conserved, indicating that natural selection conserved the encoded amino acids, for at least part of the history.

In the end, two strong candidates were found for host-gene-derived fossils predating the last common ancestor of jawed vertebrates (fig. 6). These human regions are aligned to alligator, turtle, coelacanth, and chimaera, and are not annotated as protein-coding in any of these genomes. The DNA-protein alignments have frameshifts (fig. 6 B, D), and the basewise conservation does not suggest 3-periodicity (not shown). Their ancient conservation, and strong phastCons conservation in mammals, testifies to their exaptation for some critical but unknown function.

**Fig. 6.**
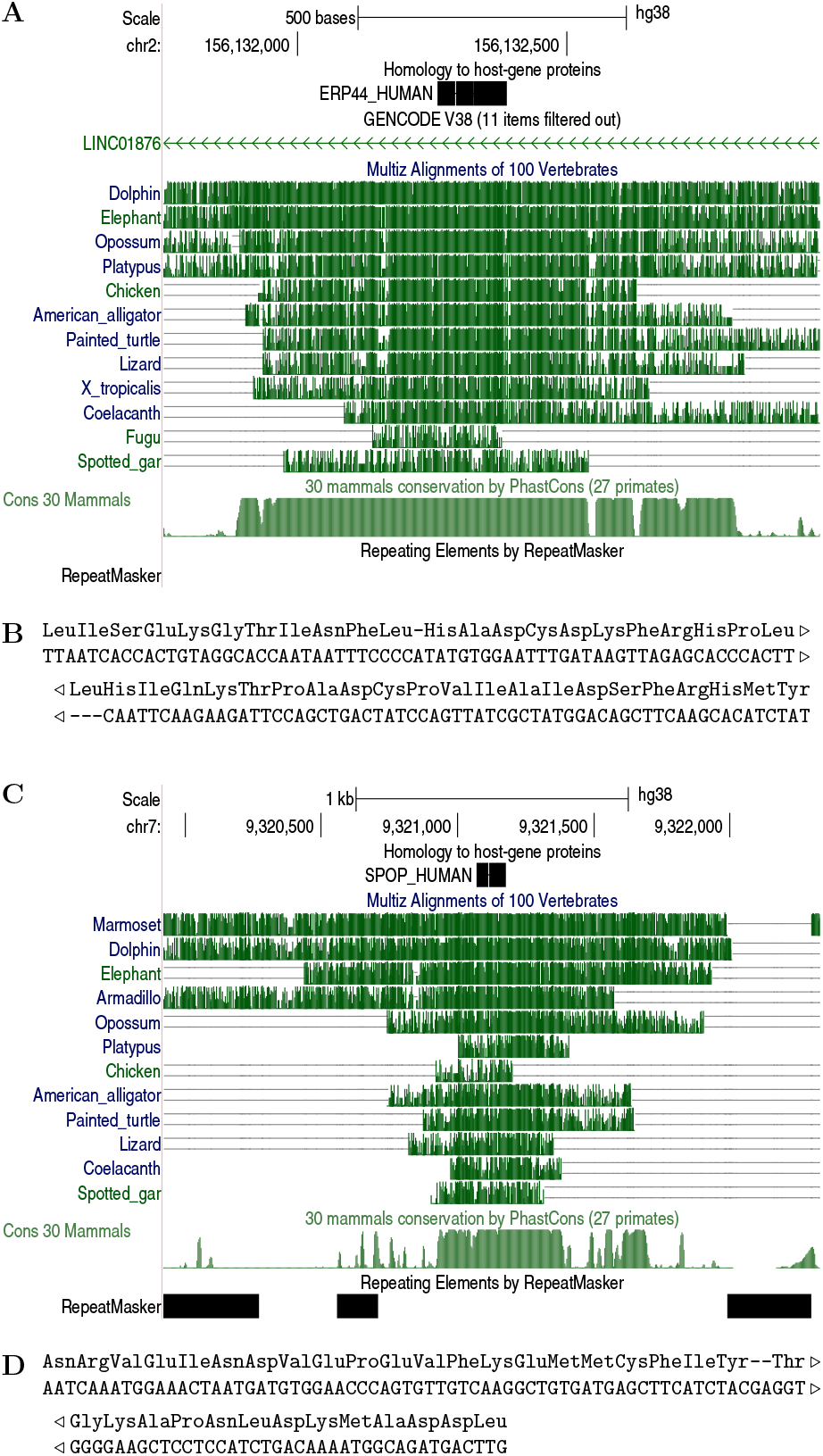
Ancient conserved pseudogenes in the human genome. **A** Match between endoplasmic reticulum resident protein 44 and chromosome 2, showing conservation in vertebrates. **B** Base-level alignment of the above. **C** Match between speckle-type POZ protein and chromosome 7. **D** Base-level alignment thereof.

### Conclusions and Prospects

This study greatly increases the number and variety of Paleozoic protein fossils. Fossils of most major TE categories (except Helitrons) are found that predate the amphibian/amniote divergence. The oldest fossils, from both TEs and host genes, predate the last common ancestor of jawed vertebrates. The detection of some TE types in ancestral genomes makes their distribution in vertebrates less patchy, suggesting that ancient vertebrates had a high diversity of TEs that were vertically inherited in some lineages but lost activity in others. There are hints that marine or aquatic vertebrates are prone to horizontal TE transfer (Ivancevic et al. 2018; Zhang et al. 2020; Barreat and Katzourakis 2021b), which might explain the high ancestral diversity. These ancient fossils have strong sequence conservation, indicating exaptation, and some have evidence of regulatory function. Not only TEs but also host-gene fossils were anciently exapted with strong sequence conservation. Ancient fossils can be markers of long-term genome tectonics.

It is hoped that these fossil-finding methods can easily be adapted for future studies. They are especially beneficial for finding TEs in less-studied genomes, reducing reliance on de novo repeat-finding and confusion between low copy-number TEs, multi-gene families, and TE-derived genes (Arkhipova 2017; Makalowski et al. 2019). The fitting of substitution and gap rates could perhaps be improved: here it was done naively by comparing a genome to known TE proteins. The choice of sequence data for parameter-fitting seems important for finding ancient or unknown types of fossil. Fossil-finding could also be aided by ancestralizing the genome sequence, e.g. reverting recent substitutions and TE insertions.

One promising application is paleovirology: few Mesozoic and no Paleozoic viral fossils have been found so far (Barreat and Katzourakis 2021a). If Gypsy-like elements (Metaviridae) or Polintons are counted as viruses, Paleozoic fossils predating ∼350 million years are found here (table 4).

A great challenge is to infer ancient genetic sequences from their fossil fragments, much as ancient organisms are inferred from mineral fossils. This inference might be assisted by LAST’s ability to estimate the probability that each column of a sequence alignment is correct.

## Materials and Methods

The pipeline scripts are available at: https://gitlab.com/mcfrith/protein-fossils.

### Genome Data

Genome sequences and their RepeatMasker annotations were downloaded from UCSC, NCBI, or repeatmasker.org (table 1). The human RepeatMasker annotations are from UCSC’s `rmskOutCurrent`.

TE protein sequences were taken from the file `RepeatPeps.lib` in RepeatMasker version 4.1.2-p1. For each non-human genome, proteins encoded by host genes were taken from NCBI’s `.faa` file for that genome. For human, with the aim of getting reliable proteins, non-TE proteins with existence level 1– 3 were taken from `uniprot sprot human.dat` in UniProt release 2021 02 (The UniProt Consortium 2020).

Protein-coding regions of the human genome were taken from the union of `wgEncodeGencodeCompV37` and `ncbiRefSeq` from UCSC (Pruitt et al. 2014; Harrow et al. 2012). For each non-human genome, protein-coding regions were obtained from NCBI’s`.gff` file for that genome.

### Finding Protein Fossils

The DNA/protein substitution and gap rates were found separately for each genome, by comparing it to the TE proteins, using LAST version 1250:

~~~
lastdb -q -c myDB RepeatPeps.lib
last-train -P8 --codon -X1 --pid=50
      myDB genome.fa > te.train
~~~

The `--pid=50` option makes it only use homologies with ≤ 50% identity, with the aim of focusing on old fossils. Next, the genome was matched to TE and host-gene proteins:

~~~
fasta-nr hostProteins RepeatPeps.lib |
    lastdb -q -c pDB
lastal -D1e9 -K0 -m500 -p te.train
        pDB genome.fa > aln.maf
~~~

Option `-D1e9` sets the significance threshold to one false hit per 10^9^ bp, `-K0` omits hits that overlap stronger hits in the genome, and `-m500` makes it more slow and sensitive. (With lower values of `m`, occasionally a host-gene-derived fossil was missed and instead wrongly aligned to a TE protein.)

It turns out the RepeatMasker proteins include exapted genes: they were excluded, by omitting hits to proteins whose names contain `_HSgene`, `_Hsa_`, `UN-GIN`, or `_Xtr_eg_tp`.

Finally, alignments >10% covered by protein-coding annotation were removed, as were host-protein alignments >10% covered by RepeatMasker TE annotations other than `Low complexity` and `Simple repeat`.

### Genome Alignments

Pair-wise genome alignments were made like this:

~~~
lastdb -P8 -uMAM8 gDBgenome1.fa
last-train -P8 --revsym -D1e9 --sample-number=5000
             gDB genome2.fa > g.train
lastal -P8 -D1e9 -m100 -p g.train gDB genome2.fa |
   last-split -fMAF+ > many-to-one.maf
last-split -r many-to-one.maf |
   last-postmask > one-to-one.maf
~~~

The `-uMAM8` and `-m100` options make it extremely slow and sensitive (Frith and Noée 2014). These one-to-one alignments are available at https://github.com/mcfrith/last-genome-alignments.

Next, isolated alignments were removed by defining two alignments to be “linked” if, in both genomes, they are separated by at most 10^6^ bp and by at most 5 other alignments. Alignments were retained if linked, directly or indirectly, to at least two others.

### Ancient Protein Fossils

A protein fossil was inferred to be ancient if it overlaps an inter-genome alignment. However, spurious overlaps are caused by the DNA-protein or intergenome alignments overshooting beyond the end of homology: this often happens when the fossil is near a protein-coding exon. Therefore, the set of alignments between two genomes was reduced to those that do not overlap protein-coding annotations in either genome, and then each fossil was considered conserved if at least 30% of it is covered by alignments between those two genomes. This 30% threshold was determined empirically (25% was occasionally insufficient). There is likely a better way using LAST’s ability to estimate the probability of each column in an alignment.

### Novelty

For tables 4 and 5, a TE fossil was deemed “known” if it has non-zero same-strand overlap with a RepeatMasker genome annotation of the same “class” (DNA, LINE, LTR, etc). For table 1, a TE fossil was deemed novel if at most 10% of it is covered by same-strand RepeatMasker TE annotations with known “class/family”. A host-gene protein fossil was deemed novel if at most 10% of it overlaps same-strand known pseudogenes.

### Nearest genes

The nearest genes were found from among those with `NM_` accession numbers in `ncbiRefSeqCurated` from UCSC.

## Acknowledgments

I thank the Frith and Asai labs’ members, past and present, for useful discussions and feedback, and Ayaka Ishiguro for a preliminary survey of protein fossils in eukaryote genomes.

